# Differences in stomatal sensitivity to CO_2_ and light influences variation in water use efficiency and leaf carbon isotope composition in two genotypes of the C_4_ plant *Zea mays*

**DOI:** 10.1101/2023.12.01.569655

**Authors:** Joseph D Crawford, Robert J. Twohey, Varsha S. Pathare, Anthony J. Studer, Asaph B. Cousins

## Abstract

The ratio of net CO_2_ uptake (*A_net_*) and stomatal conductance (*g*_s_) is an intrinsic measurement of leaf water use efficiency (*WUE*_i_) however its measurement can be challenging for large phenotypic screens. Measurements of leaf carbon isotope composition (δ^13^C_leaf_) may be a scalable tool to approximate *WUE*_i_ for screening because it in part reflects the competing influences of *A*_net_ and *g*_s_ on the CO_2_ partial pressure (*p*CO_2_) inside the leaf over time. However, in C_4_ photosynthesis the CO_2_ concentrating mechanism complicates the relationship between δ^13^C_leaf_ and *WUE*_i_. Despite this complicated relationship, several studies have shown genetic variation in δ^13^C_leaf_ across C_4_ plants. Yet there has not been a clear demonstration of whether *A*_net_ or *g*_s_ are the causal mechanisms controlling *WUE*_i_ and δ^13^C_leaf_. Our approach was to characterize leaf photosynthetic traits of two *Zea mays* recombinant inbred lines (Z007E0067 and Z007E0150) which consistently differ for δ^13^C_leaf_ even though they have minimal confounding genetic differences. We demonstrate that these two genotypes contrasted in *WUE*_i_ driven by differences in the speed of stomatal responses to changes in *p*CO_2_ and light that lead to unproductive leaf water loss. These findings provide support that differences in δ^13^C_leaf_ in closely related genotypes do reflect greater *WUE*_i_ and further suggests that differences in stomatal kinetic response to changing environmental conditions is a key target to improve *WUE*_i_.

## Introduction

Agricultural crop production requires large amounts of water (Gruère and Shigemitsu, 2021). However, the current rate of water use by crop plants may exceed water supply as climate change continues to alter precipitation patterns and decrease soil water availability (Hamdy *et al*., 2003). Increasing crop water use efficiency, through genetic improvement, can enhance or at least sustain crop production in water limited environments (Leakey *et al*., 2019). While the mechanics of how plants use water has been well studied (von Caemmerer and Baker, 2007) there is less known about the genetic control of plant water use efficiency. This is especially true for C_4_ crop species, despite their global importance in producing food calories and absolute irrigation requirements (Vadez *et al*., 2014; Leakey *et al*., 2019; Jobe *et al*., 2020; Hrozencik and Aillery, 2021).

Plant water use efficiency is in part influenced at the leaf level through the ratio of net CO_2_ (*A*_net_) uptake relative to the rate of water lost via transpiration through the stomata (Wong *et al*., 1979). Stomatal conductance (*g*_s_) is a key component of leaf-level water use efficiency because it helps regulate both the diffusion of CO_2_ into the leaf and water lost from the leaf to the surrounding air (Leakey *et al*., 2019). Changes in *g*_s_ are an important way that plants dynamically adjust to environmental signals to balance CO_2_ uptake and water loss. For example, under fluctuating light intensities that occur under natural growth conditions, low light decreases *A*_net_ and *g*_s_ to reduce unproductive leaf water transpiration (Lawson and Blatt, 2014). The speed of stomata closure has been shown to be species dependent (Ozeki *et al*., 2022), and to even vary between genotypes within species. In fact, several studies have demonstrated that the rate at which *g*_s_ responds to environmental conditions is a heritable trait that can be selected for to improve leaf-level water use efficiency in key crop species (McAusland *et al*., 2016; Pignon *et al*., 2021*a*).

The relationship between *A*_net_ and *g*_s_ provides an estimate of leaf-level intrinsic water use efficiency (*WUE*_i_) and can be measured with routine measurements of leaf gas exchange (Ellsworth and Cousins, 2016). However, these measurements are time and labor intensive and uncouples the leaf from its natural growth environment. This makes gas exchange measurements of *WUE*_i_ difficult to scale up to screen hundreds of genotypes in the field (Ellsworth and Cousins, 2016). To address this limitation in screening for *WUE*_i_, the use of leaf carbon isotope composition (δ^13^C_leaf_) has been proposed as potential proxy to more rapidly screen for variation in *WUE*_i_ (Farquhar *et al*., 1982; Farquhar, 1983; Farquhar and Richards, 1984; Rebetzke *et al*., 2002). While δ^13^C_leaf_ does not directly correspond to *WUE*_i_, it can provide a time-integrated estimate of the ratio of the CO_2_ partial pressure (*p*CO_2_) inside the leaf relative to *p*CO_2_ of the atmosphere (*C*_i_/*C*_a_), which has a strong relationship to *A*_net_/*g*_s_ and therefore *WUE*_i_. Theory suggests that *A*_net_ and *g*_s_ driven changes in *C*_i_/*C*_a_ will lead to corresponding changes in δ^13^C_leaf_ that reflects *WUE*_i_ (Farquhar and Richards, 1984). In C_3_ crop species δ^13^C_leaf_ has been successfully used to breed for increased *WUE*_i_ (Condon *et al*., 2004). However, in C_4_ plants the carbon concentrating mechanism (CCM) and their inherently low photosynthetic carbon isotope discrimination limit variation in δ^13^C_leaf_ and complicate its relationship to *WUE*_i_. For example, the leakage of CO_2_ from the CCM (e.g., leakiness, ϕ) can change δ^13^C_leaf_ independent of *WUE*_i_. Additionally, the low photosynthetic carbon isotope discrimination that occurs in response to changes in *C*_i_/*C*_a_ in C_4_ plants means that any non-photosynthetic discrimination can potentially mask the contribution of *C*_i_/*C*_a_ to δ^13^C_leaf_ (Henderson *et al*., 1992; Caemmerer *et al*., 2014). Therefore, the use δ^13^C_leaf_ as a proxy for *WUE*_i_ in C_4_ plants is more ambiguous and potentially limits its usefulness as a genetic screening tool for breeding *WUE*_i_ in these crops (Farquhar, 1983).

Despite this ambiguity, there are several examples demonstrating that δ^13^C_leaf_ in C_4_ plants does correspond to *WUE*_i_ and whole plant water use efficiency (*WUE*_plant_). In the C_4_ plant *Sorghum bicolor*, δ^13^C_leaf_ correlated with gas exchange measurements of *WUE*_i_ (Henderson *et al*., 1998) and earlier studies found repeatable genotypic variability in δ^13^C_leaf_ (Hubick *et al*., 1990). The relationship of δ^13^C_leaf_ and *WUE*_i_ has been further supported in *Setaria italica*, *Setaria viridis*, and maize (*Zea mays*) (Monneveux *et al*., 2007; Ellsworth *et al*., 2017; Twohey *et al*., 2019). Additionally, the influence of ϕ has been shown to be relatively consistent within C_4_ species (Caemmerer *et al*., 1997; Ubierna *et al*., 2013; Kromdijk *et al*., 2014) and did not respond to changes in soil water availability (Sonawane and Cousins, 2020) which suggests variation in ϕ has a negligible impact on δ^13^C_leaf_ in closely related indiviudals. The relationship of δ^13^C_leaf_ and *WUE*_i_ in C_4_ species has been further supported by screening progeny of bi-parental crosses or introgression populations that have associated variation in δ^13^C_leaf_ to specific quantitative trait loci (QTL) (Gresset *et al*., 2014; Ellsworth *et al*., 2020; Sorgini *et al*., 2021). Furthermore, Ellsworth et al., (2020) found that the approximate genomic region conferring variance in δ^13^C_leaf_ also impacted *WUE*_plant_ and showed that genotypes with stacked QTLs correlated with δ^13^C_leaf_ demonstrated a stronger relationship with *WUE*_plant_. The additive effects of several independent QTL on δ^13^C_leaf_ may explain why some studies have not found a genetic basis for δ^13^C_leaf_ in C_4_ populations (Monneveux *et al*., 2007). Those populations may have lost alleles controlling variation in δ^13^C_leaf_ through inadvertent selection or lacked allelic variation for QTL associated with δ^13^C_leaf_ (O’Leary, 1988; Tieszen and Grant, 1990; Cabrera-Bosquet *et al*., 2009; Gresset *et al*., 2014). In summary, evidence suggests a genetic control of δ^13^C_leaf_ that can be linked to *WUE*_i_ and *WUE*_plant_. However, the mechanisms related to *A*_net_ and *g*_s_ controlling the relationship of δ^13^C_leaf_ and *WUE*_i_ in closely related C_4_ genotypes has not been well demonstrated. To resolve the genetic architecture of δ^13^C_leaf_ and its relationship to *WUE*_i_ it is critical to determine drivers influencing the relationship of δ^13^C_leaf_ and *WUE*_i_ in closely related individuals but with allelic diversity associated with differences in δ^13^C_leaf_.

The *Z. mays* Nested Association Mapping (NAM) population was developed to help identify the genetic control of the phenotypic diversity through multi-parent biparental linkage mapping (Yu *et al*., 2008; McMullen *et al*., 2009). To do this, twenty-six diverse founding lines were crossed with the *Z. mays* reference genotype B73. The resulting crosses are genomic mosaics with *de novo* phenotypic diversity that can be associated with genomic regions through QTL mapping (Gage *et al*., 2020). Heritable variation in δ^13^C_leaf_ was found in the founding NAM lines (Kolbe *et al*., 2018) and recombinant inbred lines (RILs) generated from two founders with divergent δ^13^C_leaf_ (B73 and CML333; the RILs are collectively referred to as the NAM-Z007 population), have significant and consistent variation in δ^13^C_leaf_ (Twohey *et al*., 2019; Sorgini *et al*., 2021). From this NAM-Z007 population of 200 RILs two siblings, Z007E0067 and Z007E0150, were identified as having contrasting and heritable δ^13^C_leaf_ across years that was related to differences in rates of whole plant transpiration (Twohey *et al*., 2019). However, it is unknown what physiological traits drive these differences in δ^13^C_leaf_. Here we hypothesize that the variation in δ^13^C_leaf_ between Z007E0067 and Z007E0150 comes from differences in leaf photosynthetic traits. We tested this hypothesis by measuring *A_net_* and *g_s_*responses to changes in *p*CO_2_ and light availability. Our data demonstrates that Z007E0067 and Z007E0150 differ in the speed at which *g*_s_ responds to dynamic changes in *p*CO_2_ and light independent of *A*_net_ that directly influences *WUE*_i_.

## Materials and Methods

### Field and greenhouse plant growth conditions

*Zea mays* genotypes Z007E0067 and Z007E0150 were planted 15 kernels per row in a field trial in 3.7 m rows with 0.8 m spacing between rows and 0.9 m alleys. Non irrigated field trials were conducted over three years with these genotypes planted 2015-05-14, 2016-05-07, and 2017-05-16 (Twohey *et al*., 2019). Seeds were attained through the USDA Germplasm Resources Information Network (GRIN).

Subsequently, Z007E0067 and Z007E0150 were grown in the University of Illinois Plant Care Facility in Urbana, IL. Kernels were germinated in 50 cell trays in LC1 soilless media SunGro Sunshine Mix #1. Seedlings were transplanted 17 days after planting into 14.55 l pots with soilless media containing LC1, sterilized field soil, peat, and perlite in a 3:1:1:1 ratio. Plants were fertilized after transplant with Osmocote ® (11-4-17), dolomitic lime, phosphorous, magnesium sulfate and gypsum. Pagent® fungicide was applied after transplanting. These plants were well-watered and fertilized with 500 ml of 300 ppm CalMag every 7 days (Twohey *et al*., 2019).

### Plant growth chamber conditions for gas exchange measurements

The two *Z. mays* RIL genotypes Z00E0067 and Z007E0150 were also grown in a growth chamber (Bigfoot series, BioChambers Inc., Winnipeg, MB, Canada) in 3.2 L pots with commercial soil (Sunshine LC1; Sun Gro Horticulture), under atmospheric CO_2_ partial pressure (*p*CO_2_). The daily chamber photoperiod was 16 h and day/night temperatures were 31 and 22°C, respectively. The growth Photosynthetic Photon Flux Density (PPFD) of 1,000 µmol photons m^-2^ s^-1^ was used with daily stepwise increases and decreases over 2 h starting at dawn and dusk, respectively. Plants were watered as needed with Peters (20-20-20) with 2.5 g l^-1^ concentration, 2.3 g l^-1^ iron chelate fertilizer (Sprint 330, BASF, Research Triangle Park, NC, USA), and 2.5 mL STEM granular micronutrient fertilizer per 4 l water (JR Peters Inc., Allentown, PA, USA). Plants position in the chamber was scrambled at each watering.

### Plant gas exchange

Plants were grown in growth chambers for about 25 days when they were randomly ordered to be immediately measured for gas exchange. For rapid CO_2_ response curves (*A*_net_/*C*_i_) the youngest, fully expanded leaf was used, typically leaf V7. A LI-6800 photosynthesis system (LI-COR Biosciences, Lincoln, NE) equipped with a 1×3 cm leaf area cuvette was used to determine leaf net CO_2_ assimilation rates (*A*_net_) in response to *p*CO_2_ in the following order: 37.5, 0, 0.93, 2.34, 3.28, 4.68, 7.02, 10.77, 14, 18.73, 23.4, 28.1, 32.8, 37.5, 56.2, 70.24, and 37.5 Pa CO_2_ at *t_leaf_* of 30°C, 70% relative humidity, and 2,000 µmol photons m^-2^ s^-1^. Leaves were acclimated for approximately 20 minutes at 37.5 Pa CO_2_ before beginning measurements.

Gas exchange response of *A*_net_ and stomatal conductance (*g*_s_) were measured in response to step-changes in *p*CO_2_ from 37, 13.9, and 69.8 Pa using the LI-6800 time-course program. Measurements were logged every 5 sec and the LI-6800-13 cuvette (surface area of 36 cm^2^) was set to *t_leaf_* of 30°C, 70% relative humidity, and 2,000 µmol photons m^-2^ s^-1^. The program held the *p*CO_2_ for 20 minutes, progressing from 37, to 13.9, to 69.8 Pa CO_2_. The response to increases and decreases in *p*CO_2_ was fit to the sigmoidal model as described in Vialet-Chabrand et *al*. (2017*a*). Where the time constants for an increase in *g*_s_ (*k_i_*) and the initial lag time (λ) were solved for and used to calculate the time to reach 90% of the total change *A*_net_ or *g*_s_ (*t*_90%_) made between maximum and minimum for comparison and statistics.

Measurements of *A*_net_ and *g*_s_ were also made in response to step-changes in light with the LI-6800 at *t_leaf_* of 30°C, 70% relative humidity, and 37 Pa *p*CO_2_. Leaves were acclimated for 20 min and then were measured at 300 µmol photons m^-2^ s^-1^ for 15 min and subsequently the light intensity was increased to 1,800 µmol photons m^-2^ s^-1^ for 15 min and then the light was returned to 300 µmol photons m^-2^ s^-1^ for another 15 min using and autoprogram. The *t*_90%_ for the maximum and minimum were calculated and reported as for the response to CO_2_.

### On-line stable isotope measurement

The LI-6800 photosynthesis system (LI-COR Biosciences, Lincoln, NE) with a 36 cm^2^ (6×6 cm) square chamber was coupled with a tunable diode laser absorption spectroscope (TDLAS; Model TGA 200A, Campbell Scientific Inc., Logan, Utah, USA) to measure the isotope flux of CO_2_ (^12^CO_2_, ^13^CO_2_). The air flow entering the leaf chamber was partially diverted to the reference line of the TDLAS, while the airflow leaving the chamber delivered to the sample line of the TDLAS (Ubierna *et al*., 2013). For the TDLAS, the dependency of ^13^C composition (δ^13^C) on CO_2_ concentration was corrected with the concentration series method (Tazoe *et al*., 2011; Ubierna *et al*., 2013, 2017). Leaf gas exchange variables and isotopes were measured in high light (1,400 µmol m^-2^ s^-1^) and then low light (300 µmol m^-2^ s^-1^).

### Carbon isotope signature of dry leaf material

For the field grown samples, the upper most fully expanded leaf was sampled with a 0.5 cm diameter hole punch in the middle of the leaf blade from both sides of the midrib. Leaves used in gas exchange were cut and dried at 65°C for 7 days. Samples of leaf material was pulverized and homogenized with stainless steel balls for 2 min in 2 ml plastic centrifuge tubes. The powder was placed in tin capsules and combusted into N_2_ and CO_2_ in an elemental analyzer (ECS 4010, Costech Analytical, Valencia, CA). The combusted gases are separated in a 3 m gas chromatography column and the isotope signature of the gases are measured with a continuous flow isotope ratio mass spectrometer (Delta PlusXP, Thermofinnigan, Bremen). Isotope ratios are reported per mill relative to Vienna Peedee belemnite standard (VPDB). Samples were measured in the Washington State University Stable Isotope Core Laboratory.

### Modeling ***Δ***^13^C from Ci/Ca under step-changes to light

The C_4_ leaf carbon isotope discrimination (Δ^13^C) was modeled as a function of the gas exchange measured *C*_i_/*C*_a_ under step-changes presented in Figure 2. The Δ^13^C was modeled from the simplified equation in Farquhar (1983). Isotope fractionation factors were held constant for both genotypes and leakiness was assumed to be 0.2 (unitless). The fractionation of ^13^CO_2_ from diffusion in air (*a*) was 4.4‰ and the fractionation of ^13^CO_2_ by Rubisco (*b*3) was 29‰. The net fractionation of ^13^CO_2_ by hydration, dissolution, and PEPC (*b*4) was calculated for the 30°C measurement temperature as −5.2‰ (Ubierna *et al*., 2017). The fractionation effect of CO_2_ as it leaks from the bundle sheath cells (*s*) was 1.8 (‰). The potential effect of *g*_m_ on Δ^13^C was modeled as either 2 µmol m^-2^ s^-1^ Pa^-1^ to estimate *C*_m_ or *g*_m_ was assumed infinite, where *C*_i_=*C*_m_ in the follow equation:

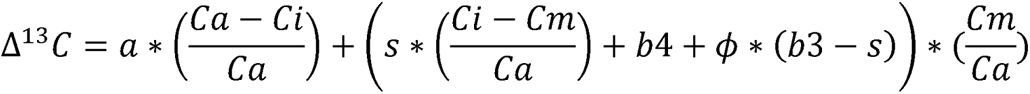

### Fluctuating Light Chamber Setup

The *Z. mays* genotypes Z00E0067 and Z007E0150 were grown together in a walk-in growth chamber (Enconair Ecological Chamber, Model GRC 36, Winnipeg, MB, Canada) under a custom built fluctuating supplemental LED system similar to Schneider et al., (2019) over a 16 hour photoperiod. The 1,500 W LED fluctuating lights (HIGROW LED, Model GLB1500, Shenzhen China) were attached to a shelf inside the walk-in chamber with the fluctuating high light (FHL), high constant light (HCL), and the low constant light (LCL) treatments. The FHL treatment used a timer to cycle the supplemental LEDs on for 5 min and then off for 7 min throughout the photoperiod. The HCL used an identical LED lamp but was on continuously throughout the photoperiod. The LCL treatment was on a lower shelf and received diffuse light from the chamber and two overhead fluorescent tube lamps. Nine biological replicates per genotype were simultaneously grown under the FHL and HCL treatments with one plant per pot that was randomly shuffled under their growth light treatment. The LCL shelf had accommodated 18 replicates of each genotype (Schneider *et al*., 2019).

### Statistical Analysis

Statistical differences between the two genotypes were assessed using one-way ANOVA with the aov function in R 3.6.3 (R Core Team, 2018). Stomatal response times were statistically assessed by, modeling k_i_ (time constants for increase/decrease), λ (a time delay), G_s-min_, and G_s-max_ to calculate *t*_90%gs_, and then performing a one-way ANOVA on *t*_90%gs_ (Vialet-Chabrand *et al*., 2017*a*). For experiments with a treatment like light level or *p*CO_2_ a two-way ANOVA was used and post-hoc tests were corrected for with Tukey’s pairwise test. For the fluctuating light experiment a two-way ANOVA was used with Tukey’s test according to the model: Y ∼ LightTreatment + Genotype + (LightTreatment*Genotype) + Error. For online stable isotope measurements extreme data points whose absolute value was 1.5 times larger than the interquartile range were removed.

## Results

### The leaf carbon isotope composition (***δ***^13^C_leaf_) and stomatal traits in selected *Zea mays* Nested Association Mapping RILs

Survey of δ^13^C_leaf_ in the *Z. mays* Nested Association Mapping population NAM-Z007 identified two lines (Z007E0067 and Z007E0150) that differed in δ^13^C_leaf_ during 2015, 2016, 2018, and 2019 field trials as well as in a greenhouse grow out (Table 1). These differences in δ^13^C_leaf_ between Z007E0067 and Z007E0150 was further replicated in growth chamber plants used for the gas exchange measurements presented in this study (Table 1). Measurements during the 2019 grow out, showed that the two genotypes were morphologically similar and did not have significant differences in stomata density on either abaxial or adaxial surface, stomatal ratio between the two surfaces, or the specific leaf area (Table 1). There was also no statistical difference in these stomatal traits between the genotypes used for the gas exchange measurements presented in the current study (Table 1). The stomatal width was not statistically different between genotypes, but stomatal length was approximately 4 µm longer on average in Z007E0150 compared to Z007E0067 (Table 1).

**Table 1.** Phenotypic characterization of full siblings of *Zea mays* genotypes Z007E0067 and Z007E0150 with contrasting for δ^13^Cleaf.

|  | Genotype |  | Statistics |
| --- | --- | --- | --- |
|  | Z007E0067 | Z007E0150 |  |
| $\delta^{13}\text{C}_{\text{leaf}}$ WSU growth chamber (‰)<sup>#</sup> | -13.2±0.1 | -14.2±0.1 | * |
| $\delta^{13}\text{C}_{\text{leaf}}$ Field Grown Plants (‰) | | | |
| 2015 (Sorgini *et al.*, 2021) | -11.64 | -13.35 | * |
| 2016 (Twohey *et al.*, 2019) | -11.85 | -13.03 | - |
| 2018 | -12.05 | -12.27 | NS |
| 2019 | -12.08 | -12.41 | * |
| Illinois greenhouse (Twohey *et al.*, 2019) | -13.15 | -13.77 | * |
| 2019 leaf anatomical data Illinois field |  |  |  |
| Stomatal Density Abaxial (stomata 0.64mm<sup>-2</sup>) | 62.8±3.0 | 61.0±3.0 | NS |
| Stomatal Density Adaxial (stomata 0.64mm<sup>-2</sup>) | 48.1±3.0 | 45.0±3.0 | NS |
| Leaf Area (cm<sup>2</sup>) | 588±34 | 509±21 | NS |
| Specific Leaf Area (cm<sup>2</sup> g<sup>-1</sup>) | 196±9 | 201±8 | NS |
| 2020 leaf anatomical data WSU growth chamber<sup>#</sup> |  |  |  |
| Stomatal Density Abaxial (stomata 1.2 mm<sup>-2</sup>) | 96.5±4.4 | 88.3±3.8 | NS |
| Stomatal Density Adaxial (stomata 1.2 mm<sup>-2</sup>) | 61.8±3.5 | 64.0±4.3 | NS |
| Total stomatal density (stomata 1.2 mm<sup>-2</sup>) | 158±7 | 152±6.3 | NS |
| Stomatal length (μm) | 43.6±2.0 | 47.8±2.3 | * |
| Stomatal width (μm) | 27.0±1.8 | 26.0±1.4 | NS |

<sup>#</sup> $\delta^{13}\text{C}_{\text{leaf}}$ and stomatal data for plants used in gas exchanged measurements presented here. \*Indicates statistically different between genotypes from a T-test at $p < 0.05$ . NS – not significant

### Gas exchange responses to rapid changes in CO_2_ availability

We measured the rates of leaf net CO_2_ assimilation (*A*_net_) and stomatal conductance (g_s_) in response to rapid (<1 min) changes in the CO_2_ partial pressures surrounding the leaf (*C*_a_). The inter cellular *p*CO_2_ (*C*_i_) increased with *C*_a_ in both genotypes but had a lower slope in Z007E0150 compared to Z007E0067 (Fig. S1). There was not a significant difference in the initial slope of *A*_net_ plotted against *C*_i_ (*A*_net_/*C*_i_) or the the saturating rate of *A*_net_ at high *C*_i_ between the two genotypes (Fig. 1A). For both genotypes, *g*_s_ declined from the lowest to highest *C*_a_; however, the *g*_s_ in Z007E0150 declined by 81%, compared to a 61% decline in Z007E0067 (Fig. 1B). These differences in *g*_s_ lead to different relationship of *A*_net_ with *g*_s_, where the intrinsic leaf water use efficiency (*WUE*_i_; *A*_net_/*g*_s_) was greater during these short-term changes in *p*CO_2_ in Z007E0150 compared to Z007E0067 (Fig. 1C).

**Figure 1.**
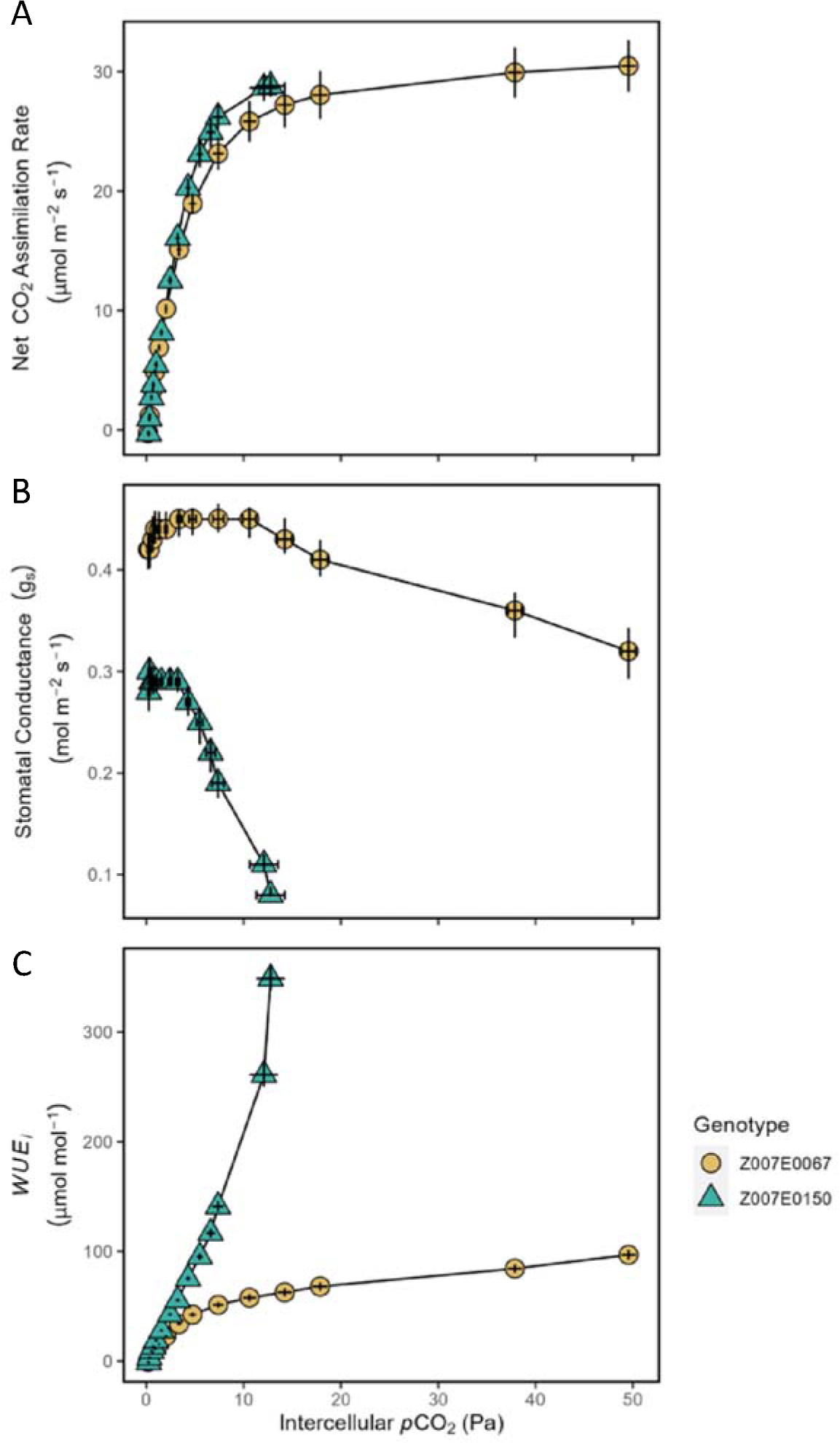
The net CO_2_ assimilation rate (*A*_net_), stomatal conductance (*g*_s_), and intrinsic water use efficiency (*WUE*_i_; calculated as *A*_net_/*g*_s_) response to intercellular *p*CO_2_ (Pa) in the *Z. mays* genotypes Z007E0067 and Z007E0150 with contrasting carbon isotope composition (δ^13^C_leaf_). The measurements were at a leaf temperature of 30°C, 70% relative humidity, and Photosynthetic Photon Flux Density (PPFD) of 1,800 µmol m^-2^ s^-1^. Symbols in all panels are mean value ± SE (n=6).

### Response to step-changes in steady-state CO_2_ availaibility

To further examine the response of *A*_net_ and *g*_s_ to changes in CO_2_, leaves were acclimated at ambient *p*CO_2_ (37 Pa) for 20 minutes and then subjected to a step-change to low *p*CO_2_ (13.9 Pa) for 20 minutes and a subsequent high *p*CO_2_ (69.8 Pa) for 20 minutes. The *A*_net_ responded rapidly to changes in *p*CO_2_ similarly between genotypes (Fig 2A). Generally, *g*_s_ was higher in Z007E0067 than in Z007E0150 under all *p*CO_2_ conditions (Fig 2B) and the genotypes also contrasted in the speed of stomatal responses to change in *p*CO_2_. During the transition to low *p*CO_2_, Z007E0150 was ca. 3-times faster to reach *t*_90%gs_ opening than Z007E0067 (Table 2; *p* < 0.01). Likewise, in response to high *p*CO_2_ the rate of *t*_90%gs_ closure was ca. 4-times faster in Z007E0150 than in Z007E0067 (Table 2; *p* < 0.01). The Z007E0067 was 10 min slower than Z007E0150 to reach the same minimum *g*_s_ under the high *p*CO_2_ (Fig. 2B). These differences in *A*_net_ versus *g*_s_ led to higher *WUE*_i_ in Z007E0150 compared to Z007E0067, particularly under ambient and elevated *p*CO_2_ (Fig. 2C).

**Figure 2.**
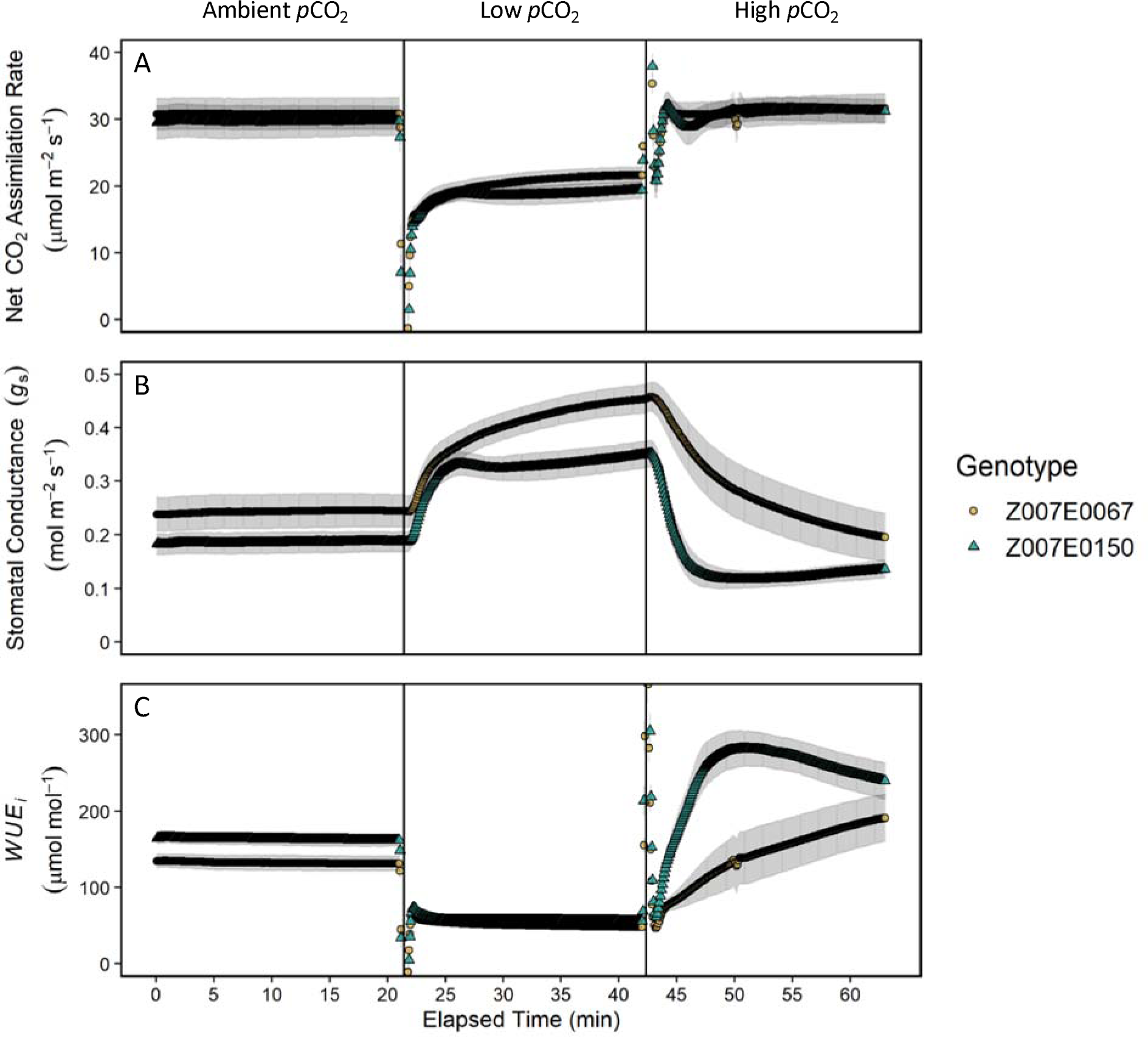
A time course response of net CO_2_ assimilation rate (*A*_net_) stomatal conductance (*g*_s_) and intrinsic water use efficiency (*WUE*_i_; calculated as *A*_net_/*g*_s_) to step-changes in *p*CO_2_ in the *Z. mays* genotypes Z007E0067 and Z007E0150 with contrasting carbon isotope composition (δ^13^C_leaf_). The step-changes in *p*CO_2_ went from ambient (37 Pa) to low (13.9 Pa) and then to high (69.8 Pa). Each period of *p*CO_2_ lasted about 20 min. Data was collected on n=6 (Z007E0067) and n=5 (Z007E0150) and data points were collected every 5 sec. Symbols in all panels are mean value ± SE.

**Table 2.**
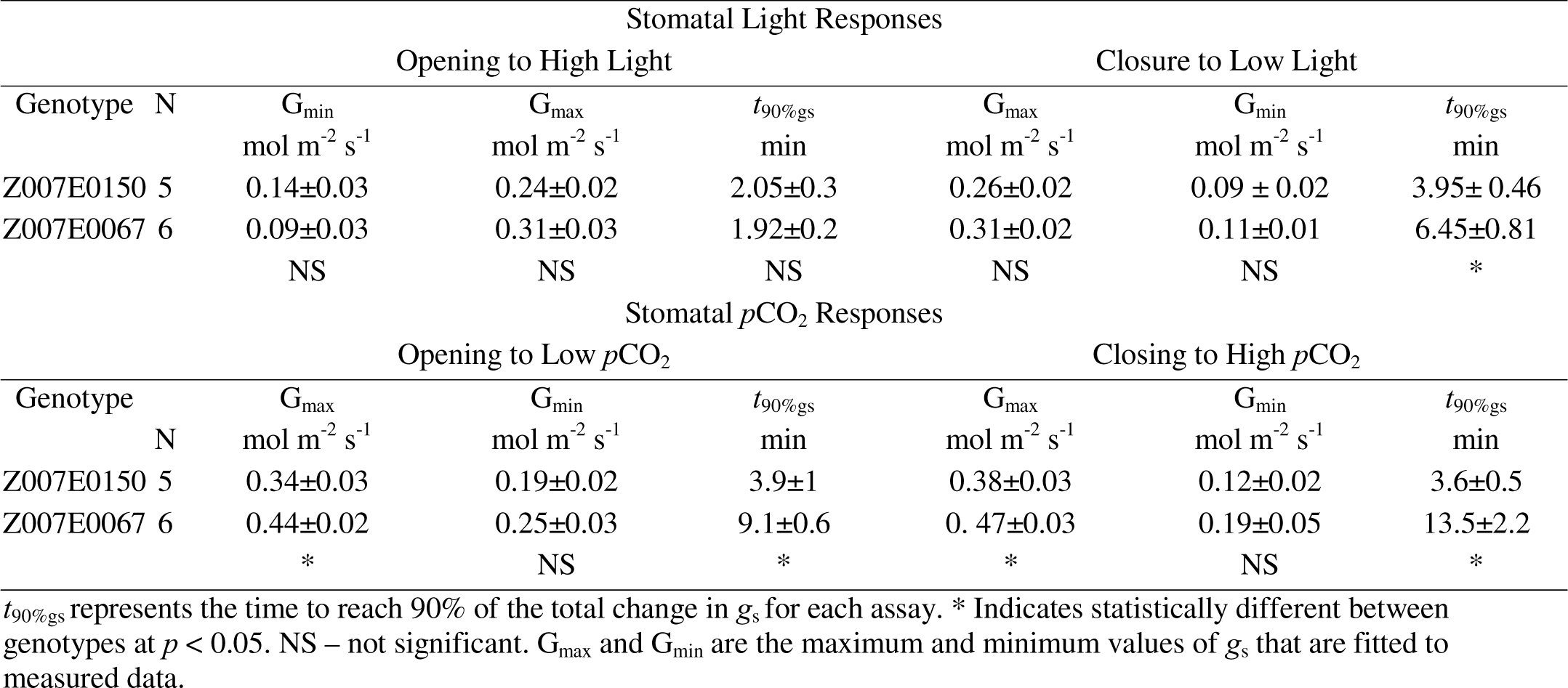
Stomatal responses to light and CO_2_.

### Response to step-changes in steady-state light conditions

Measurements of *A*_net_ and *g*_s_ were also made under step-changes of PPFD 300 to 1,800 and back to 300 µmol photons m^-2^ s^-1^ under ambient *p*CO_2_ (Fig 3). Under the initial low light conditions the genotypes had similar *A*_net_ but differed in steady-state *g*_s_ and *WUE*_i_ (Fig. 3B). During the transition to high light the response time of *A*_net_ or *g*_s_ to 90% of the maximum value (*t*_90%_) was the same between genotypes (Table 2). Under steady-state high light *A*_net_ and *g*_s_ were greater in Z007E0067 compared to Z007E0150 but *WUE*_i_ was not statistically different (Fig 3). During the transitioning from high to low light the time to reach 90% of the minimum *A*_net_ (*t*_90%Anet_) was rapid and did not differ between genotypes (Table 2). However, there was a 2.5 min difference in *t*_90%gs_ closure, with Z007E0150 being 48% faster than Z007E0067 (Table 2, *p* < 0.05). These differences in the response of *A*_net_ and *g*_s_ to the step-changes in light led to higher *WUE*_i_ in Z007E0150 compared to Z007E0067 (Fig. 3C).

**Figure 3.**
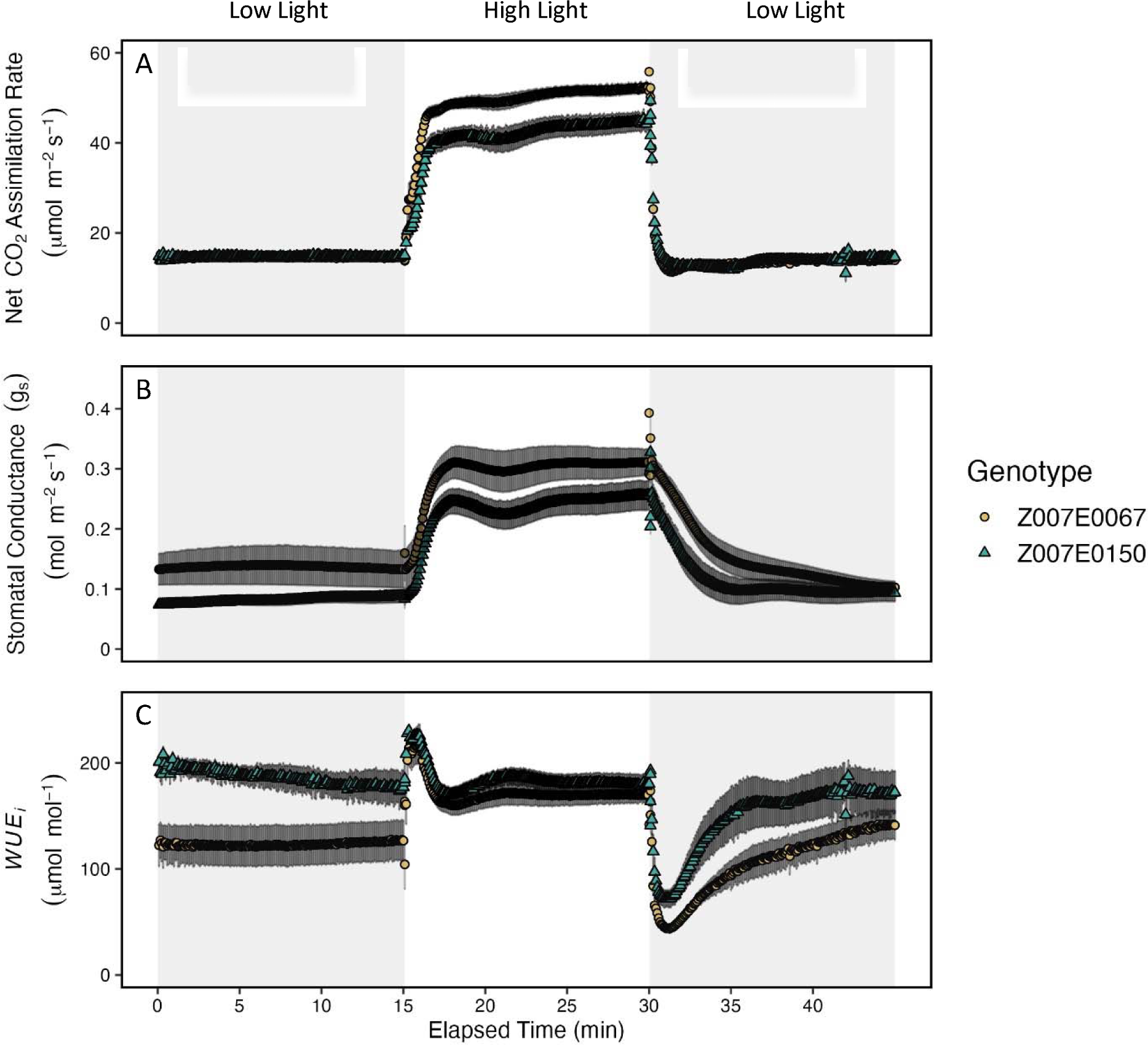
A time course response of net CO_2_ assimilation (*A*_net_), stomatal conductance (*g*_s_) and intrinsic water use efficiency (*WUE*_i_) to step-changes in light intensity in the *Z. mays* genotypes Z007E0067 and Z007E0150 with contrasting carbon isotope composition (δ^13^C_leaf_). The low light intensity was 300 PPFD µmol m^-2^ s^-1^ followed by a step change to 1,800 PPFD µmol m^-2^ s^-1^ for 15 minutes, and then returned to 300 PPFD µmol m^-2^ s^-1^ for 15 minutes. Data was collected on n=5 (Z007E0067) and n=6 (Z007E0150) every 5 sec. Symbols in all panels are mean value ± SE.

### Comparisons of leaf carbon isotope discrimination (***Δ***^13^*C*)

Steady-state gas exchange and tunable diode laser absorption spectroscope measurements were used to simultaneously measure the ratio of intercellular to atmospheric CO_2_ partial pressure (*C*_i_/C*_a_*) and leaf isotope discrimination of ^13^CO_2_ (Δ^13^C) under steady-state conditions (Table 3). These measurements were used to test if the differences in δ^13^C_leaf_ was driven by differences between genotypes in intrinsic leaf isotope fractionation. Measurements were made at two PPFD intensities (300 and 1,400 μmol m^-2^ s^-1^) at a constant *p*CO_2_ of 37 Pa and at two *p*CO_2_ (37 and 69.8 Pa) at 1,800 PPFD (μmol m^-2^ s^-1^). The *C*_i_/C*_a_* was generally lower in the Z007E0067 than in the Z007E0150 under both light conditions and differed between measurement light conditions for both genotypes (Table 3). The *C*_i_/C*_a_* did not differ between the two *p*CO_2_ measurement conditions for either genotype (Table 3). The Δ^13^C did not differ between any of the measurement light or *p*CO_2_ conditions nor between genotype (Table 3).

**Table 3.** Comparison of steady-state ratio of intercellular to atmospheric CO_2_ partial pressure (*C*_i_/Ca) and leaf isotope discrimination of ^13^CO_2_ (Δ^13^C) between the Z007E0150 and Z007E0067 lines. Low and high light measurements were made under 300 and 1,400 PAR µmol m^-2^ s^-1^ at 37 Pa *p*CO_2_. The low and high *p*CO_2_ measurements were made under 37 and 69.8 *p*CO_2_ (Pa) and 1,800 µmol m^-2^ s^-1^. Data represents the mean value ± SE where n = 5-6. Different letter represent significant differences between genotypes within a given measurement condition.

| | $C_i/C_a$ | | $\Delta^{13}\text{C}$ (‰) | |
| --- | --- | --- | --- | --- |
|  | Z007E0150 | Z007E0067 | Z007E0150 | Z007E0067 |
| Light response ($\mu\text{mol m}^{-2} \text{s}^{-1}$) | | | | |
| 300 | 0.20±0.03<sup>a</sup> | 0.17±0.05<sup>c</sup> | 5.1±3.0<sup>a</sup> | 5.0±3.5<sup>a</sup> |
| 1400 | 0.32±0.02<sup>b</sup> | 0.26±0.02<sup>d</sup> | 4.8±1.8<sup>a</sup> | 4.2±3.1<sup>a</sup> |
| CO<sub>2</sub> (Pa) |  |  |  |  |
| 37 | 0.3±0.1<sup>a</sup> | 0.4±0.1<sup>a</sup> | 5.0±1.0<sup>a</sup> | 4.2±0.8<sup>a</sup> |
| 69.8 | 0.4±0.1<sup>a</sup> | 0.6±0.1<sup>a</sup> | 5.1±1.3<sup>a</sup> | 3.6±0.9<sup>a</sup> |

### Modeled response of **Δ**^13^C to step-changes in light and pCO_2_

Measurements of Δ^13^C under transient conditions are noisy and difficult to make in C_4_ plants. Therefore, Δ^13^C was modeled (Δ^13^C_mod_) with the measured changes in *C*_i_/*C*_a_ derived from the data determined from the step-changes in *p*CO_2_ and light (Figs 2 and 3, respectively). The *C*_i_/*C*_a_ measured from the step-changes in light (Fig. 4A) differed under the initial low light (Fig 4A) which translated to a difference in Δ^13^C_mod_ of approximately 1 ‰ (Fig 4B). After the transition to high light there was no significant difference in *C*_i_/*C*_a_ or Δ^13^C_mod_ between genotypes despite the differences in A_net_ (Fig 3A). After the initial transition back to low, the *C*_i_/*C*_a_ was higher and Δ^13^C_mod_ lower in the Z007E0067 than in the Z007E0150; however, this difference diminished towards the end of these measurement conditions (Fig. 4B). A similar trend of *C*_i_/*C*_a_ being higher and Δ^13^C_mod_ lower was seen in the Z007E0067 compared to the Z007E0150 plant when using the *p*CO_2_ response data presented in Figure 2 (Fig. S2).

**Figure 4.**
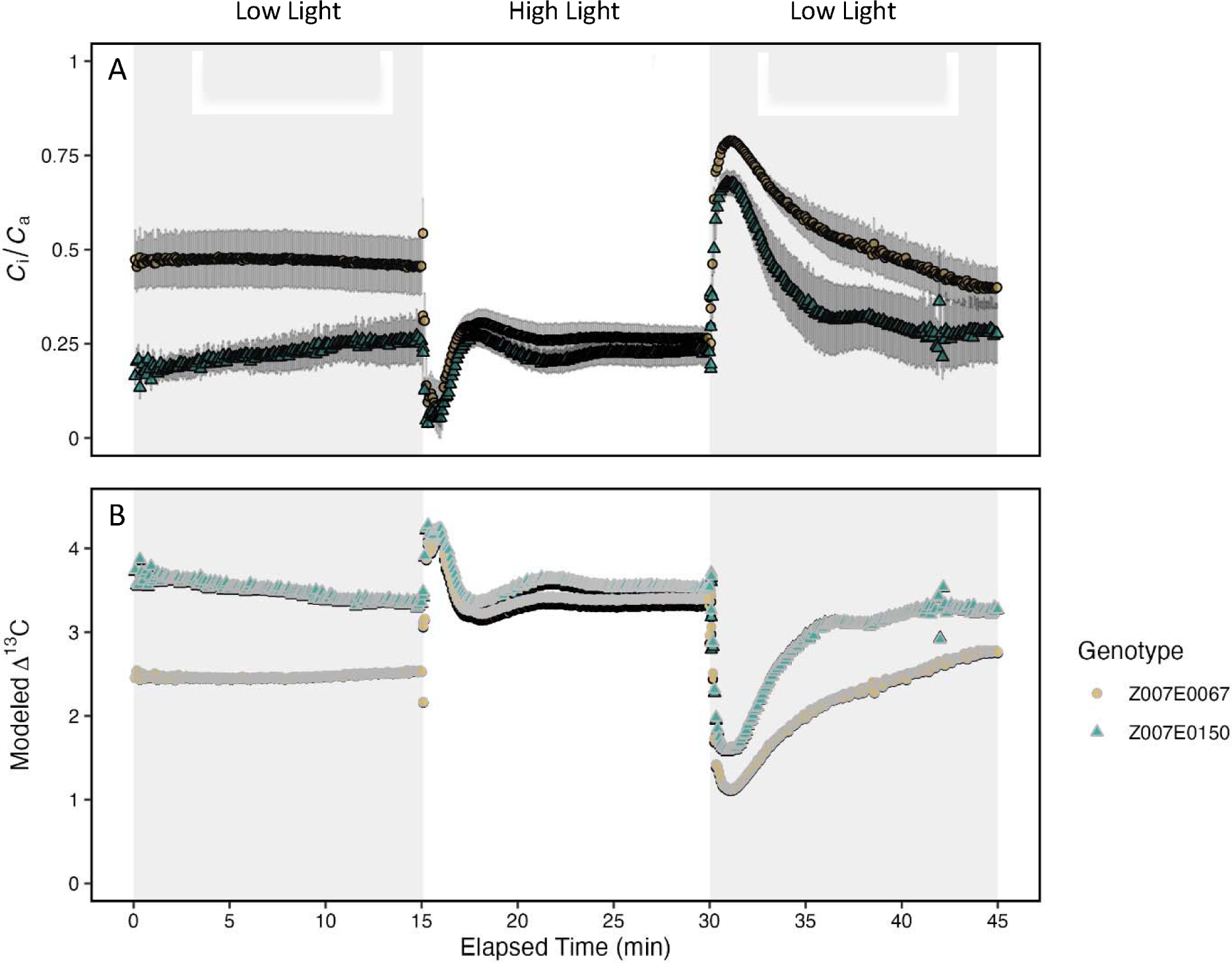
A time course response of the ratio of intercellular to atmospheric CO_2_ partial pressure and modeled Δ^13^C_mod_ to step-changes in light in two *Z. mays* genotypes contrasting for carbon isotope composition. The light conditions are described in Figure 3 as 300 PPFD µmol m^-2^ s^-1^ followed by a step change to 1,800 PPFD µmol m^-2^ s^-1^ for 15 minutes, and then returned to 300 PPFD µmol m^-2^ s^-1^ for 15 minutes. The Δ^13^C_mod_ was modeled from data collected on n=5 (Z007E0067) and n=6 (Z007E0150) and data points were collected every 5 sec.

### Growth response to fluctuating light conditions

The Z007E0150 and Z007E0067 genotypes were grown side-by-side under either Low Constant Light (LCL), High Constant Light (HCL), and Fluctuating High Light (FHL). Under these growth conditions the leaf carbon isotope composition (δ^13^C_leaf_) was significantly affected by the light treatment (Fig. 5; *p* < 0.01) but was more depleted in Z007E0150 than in Z007E0067 line under all conditions (Fig. 5; *p* < 0.01). However, the difference in δ^13^C_leaf_ between genotypes was smaller under the FHL compared to the LCL and HCL treatments.

**Figure 5.**
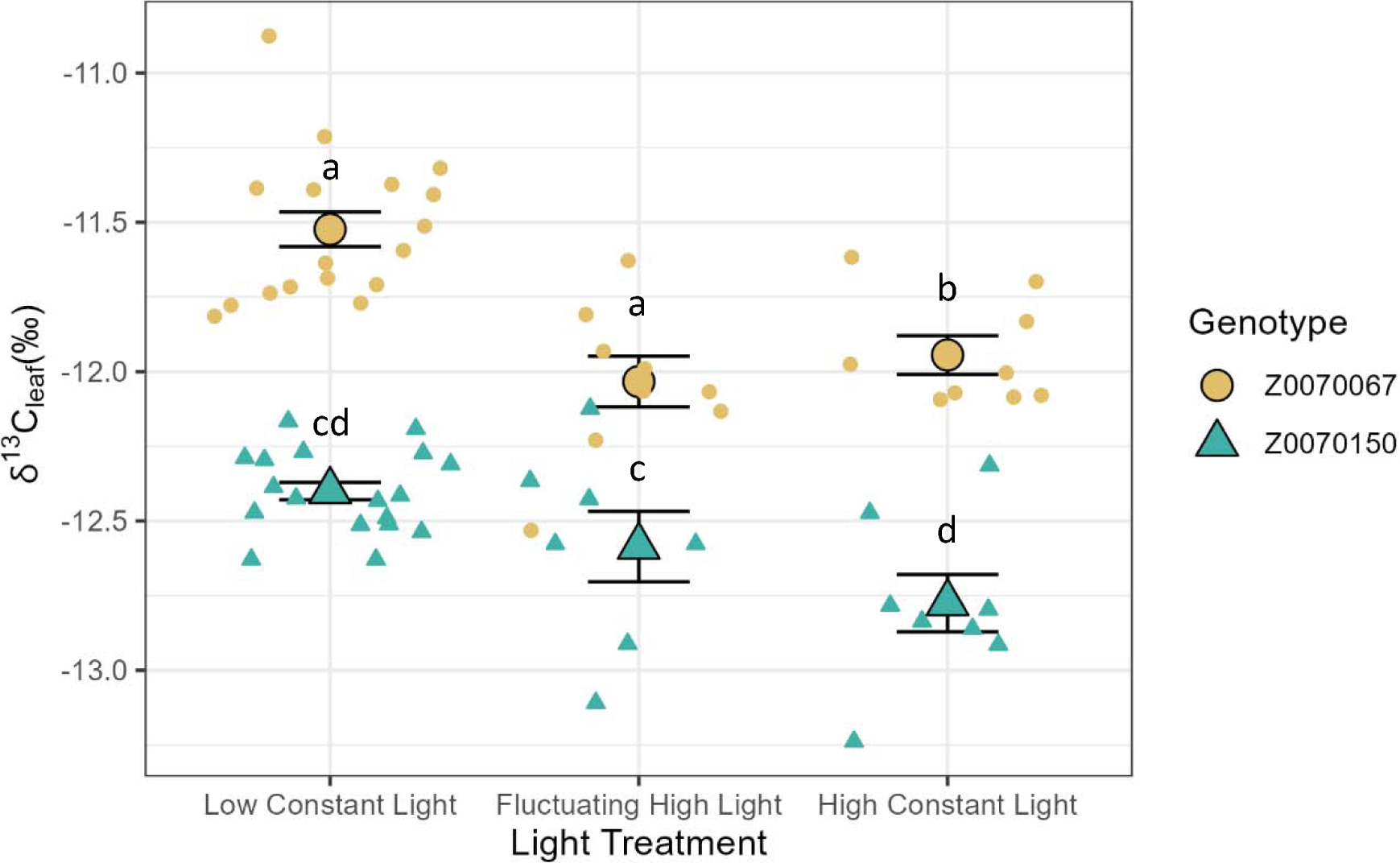
Two *Z. mays* genotypes contrasting for carbon isotope composition grown under three light treatments: Low Constant Light (LCL), Fluctuating High Light (FHL), and High Constant Light (HCL).The leaf carbon isotope composition of plants grown under the light treatments. The LED lights were inside a growth chamber that supplied ambient lighting. The fluctuating high light (FHL) alternated during the daily photoperiod between “on” (5 min where the light level approximately matched the HCL level), and then “off” where the only light was low ambient, diffuse light of the chamber. The HCL light level was from the ambient light of the chamber and the LED lamps that were on for the entire photoperiod. The LCL treatment was the diffuse lighting of the chamber but plants were on a shelf below the other treatments and heavily shaded.

## Discussion

Leaf-level intrinsic water use efficiency (*WUE*_i_) in C_4_ plants can be driven by both the net rate of CO_2_ assimilation (*A*_net_) and stomatal conductance (*g*_s_) but these traits are difficult to screen in large breeding populations. It has been well documented in C_4_ plants that *A*_net_/*g*_s_ influences the operational level of CO_2_ inside the leaf (*C*_i_/*C*_a_) and that leaf carbon isotope composition (δ^13^C_leaf_) is influence by *C*_i_/*C*_a_. This suggests that *A*_net_/*g*_s_ should be associated with heritable variation in δ^13^C_leaf_ because of their relationships with *C*_i_/*C*_a_. However, more research is needed to determine if δ^13^C_leaf_ is an effective proxy for *A*_net_/*g*_s_ in C_4_ plants because of the environmental, genetic, and other plant physiology parameters that can also influence δ^13^C_leaf_. The goal of this paper was to determine the physiological drivers of δ^13^C_leaf_ in two *Zea mays* lines with low genetic diversity but consistent and statistically significant differences in δ^13^C_leaf_.

### Identification of Zea mays siblings with distinct ***δ***^13^C_leaf_ phenotypes

Previous work has shown significant variation in δ^13^C_leaf_ across the 26 founder lines of the *Z. mays* Nested Association Mapping (NAM) population (Kolbe *et al*., 2018; Twohey *et al*., 2019). Several recombinant inbred (RIL) families have been generated from these NAM founders (McMullen et al. 2009), including RIL populations from parental lines with divergent δ^13^C_leaf_ (e.g., B73 and CML333). The RILs generated from the B73 and CML333 cross, collectively referred to as the NAM-Z007 population, was shown to have significant and consistent variation in δ^13^C_leaf_ (Twohey *et al*., 2019; Sorgini *et al*., 2021). From this population two siblings, Z007E0067 and Z007E0150, were identified among the 200 RILs as having contrasting and heritable δ^13^C_leaf_ across years that was related to differences in whole plant transpiration (Twohey *et al*., 2019). Our further analysis of the Z007E0067 and Z007E0150 siblings showed similar δ^13^C_leaf_ differences across four years of independent field trials as well as when they were grown in a greenhouse and growth chamber (Table 1). These consistent δ^13^C_leaf_ differences in Z007E0067 and Z007E0150 provide an opportunity to test the relationship of δ^13^C_leaf_ and *WUE*_i_ in two closely related *Z. mays* lines.

### The relationship of ***δ***^13^C_leaf_ and WUE*_i_*

Previous studies have shown that variation in *WUE*_i_ can occur through changes in both *A*_net_ and *g*_s_ (McAusland *et al*., 2016; Leakey *et al*., 2019; Pignon *et al*., 2021*b*; Ozeki *et al*., 2022). The measurements of *A*_net_ versus *C*_i_ (*A*_net_/*C*_i_ curves) showed no evidence that Z007E0067 and Z007E0150 differed in the initial slope of an *A*_net_/*C*_i_ curve (a proxy for the efficiency of the C_4_ CCM) or net CO_2_ assimilation under saturating *p*CO_2_ (Fig 1A). This demonstrates there was little difference in the photosynthetic capacity between these two lines but the differences in δ^13^C_leaf_ between Z007E0067 and Z007E0150 could be associated with differences in the availability of CO_2_within the leaf. In fact, the short-term measurements of *A*_net_/*C*_i_ curves show that Z007E0150 maintained a lower *C*_i_ at all measurement *p*CO_2_ due to lower *g*_s_ that was significantly more sensitive to the changes in CO_2_ availability. The lower *g*_s_ and its rapid closure in response to *p*CO_2_ lead to significantly higher *WUE*_i_ in Z007E0150 than in Z007E0067 during these measurements (Fig 1). These non-steady-state changes in *WUE*_i_ correlated with the more negative δ^13^C_leaf_ in the Z007E0150 compared to Z007E0067 suggesting that under growth conditions Z007E0150 operates at a lower *C_i_/C_a_* and increased *WUE*_i_.

The relationship between *A*_net_ and *g*_s_ is often perturbed under non-steady conditions such as the rapid changes in *p*CO_2_ used during *A*_net_/*C*_i_ curve measurement (Caemmerer, 2000). Therefore, the response of *A*_net_ and *g*_s_ was determined during a continuous time course changing from ambient, low, and high *p*CO_2_ (37, 13.9, and 69.8 Pa, respectively). The *g*_s_ in Z007E0150 compared to Z007E0067 was more than two times faster in opening and closing to low and high *p*CO_2_, respectively (Table 2). These quicker *g*_s_ responses in Z007E0150 also translated to generally higher *WUE*_i_ during elevated *p*CO_2_ conditions that invoked stomatal closure (Fig 2). Under the steady-state conditions, *A*_net_ was the same across all three measurement *p*CO_2_ confirming the photosynthetic capacity was not different between the two genotypes. However, under the state-state conditions *g*_s_ in Z007E0150 was consistently lower than in the Z007E0067 line. These differences lead to a generally higher *WUE*_i_ in Z007E0150 under the ambient and high *p*CO_2_ (37 and 69.8 Pa, respectively).

The contrast in speed of stomata closure between the two lines was also evident under changing light conditions. We invoked stomatal closure via step-changes in light in a gas exchange system and compared the speed of stomata closure. This assay approximates the naturally dynamic light intensity experienced throughout a day in a leaf because of random factors such as cloud cover or wind changes in leaf angle (Vialet-Chabrand *et al*., 2017*b*). The genotypes had no statistically significant differences in their stomatal opening times, but they had considerable differences in stomatal closing times (Table 2). Upon a decrease in light intensity, Z007E0150 decreased stomatal conductance 1.6 times faster than Z007E0067 (Fig 3; Table 2), differing by about 2.5 min. During the drop in light intensity the genotypes did not differ in *A*_net,_ but because stomata remained open longer in Z007E0067 without productive photosynthesis this lowered *WUE*_i_ (Fig 3). A plant with rapidly closing stomata during the decrease in light conserves leaf water and generally has higher *WUE*_i_ (Lawson and Blatt, 2014; Leakey *et al*., 2019). Z007E0150 seems to tightly regulate *C*_i_/*C*_a_ inside the leaf with rapid stomatal responses that minimize unproductive water loss in response to dynamic light conditions. These data suggest differences in the speed of stomatal response between these two genotypes impacts *WUE*_i_ and therefore δ^13^C_leaf_.

### Stomatal sensitivity and leaf carbon isotope discrimination

Under steady-state conditions, the measured Δ^13^C for Z007E0150 and Z007E0067 lines were not statistically different between genotypes regardless of measurement light or *p*CO_2_. The lack of variation in measured Δ^13^C demonstrates differences in δ^13^C_leaf_ between genotypes was not caused by large differences in leakiness (ϕ) or mesophyll conductance to CO_2_ (*g*_m_). The direct measurements of Δ^13^C are difficult to conduct under non-steady condition because the change in Δ^13^C in C_4_ plants is relatively small and subject to relatively large error as the measurement system transition between conditions. However, theory predicts that the influence of *A*_net_/*g*_s_ on *C*_i_/*C*_a_ can drive changes in δ^13^C_leaf_ (Farquhar, 1983) and the relationship between *A*_net_/*g*_s_, *C*_i_/*C*_a_, and δ^13^C_leaf_ has been described by mathematical models of steady-state leaf ^13^CO_2_ isotope exchange (Δ^13^C; Farquhar, 1983). Using the simplified C_4_ model of Δ^13^C (Caemmerer *et al*., 2014 See Equation 1) we modeled Δ^13^C with our measured gas exchange data (*A*_net_, *C*_i_, and *C*_a_) in response to step-changes in light and *p*CO_2_ (see Δ^13^C_mod_ in Fig 4 and SFig. 2 from data presented in Figs. 2 and 3). This shows that the lower *C*_i_/*C*_a_ in Z007E0150 compared to Z007E0067 led to a greater Δ^13^C_mod_, assuming that leakiness is similar between genotypes. At low light, the difference in modeled Δ^13^C_mod_ was about 1‰, which is close to the average observed independently in several growth conditions (Table 1). Overall, these differences in Δ^13^C_mod_ support the more depleted δ^13^C_leaf_ in Z007E0150 compared to Z007E0067 suggesting that the differences in *A*_net_/*g*_s_ that led to variation in *C*_i_/*C*_a_ are driving changes in δ^13^C_leaf_ through Δ^13^C. Taken together, the measured Δ^13^C and modeled Δ^13^C_mod_ demonstrates there is likely little intrinsic difference in photosynthetic CO_2_ isotope discrimination between the Z007E0150 and Z007E0067 lines but rather the difference in δ^13^C is related to their differences in A_net_/*g*_s_ and the impact on *C*_i_/C*_a_*. This variation in δ^13^C between genotypes was maintained under the Low Constant Light (LCL), High Constant Light (HCL), and Fluctuating High Light (FHL) growth conditions. This further demonstrates the differences in δ^13^C is maintained across different light environments and must be driven by robust trait differences between these genotypes.

### The relationship of WUE_i_ with stomatal size and patterns

The rates of *g*_s_ responses to environmental conditions are primarily dependent on anatomical and biochemical traits (Taylor *et al*., 2012). Theory and empirical data has shown that maximum *g*_s_ (*g*_smax_) is a function of stomatal pore sizes (e.g. length times width) and density per unit leaf surface area (Franks *et al*., 2009). Stomatal pore sizes and density have been correlated with *WUE*_i_ in C_4_ plants (Ferguson *et al*., 2021; Xie *et al*., 2021). These traits have also been demonstrated to influence the rates of *g*_s_ responses to environmental conditions, where smaller stomata generally respond faster to environmental stimuli than larger ones (Taylor *et al*., 2012; Drake *et al*., 2013; Lawson and Blatt, 2014; Lawson and Vialet-Chabrand, 2019). Smaller stomata are thought to increase the volume-to-surface area ratio of pores and therefore ion fluxes can increase changes in turgor more potently during transient responses (Hetherington and Woodward, 2003; Lawson and Blatt, 2014). Decreased guard cell length alone has been shown to increase stomatal response speed while holding stomata density constant but under a 16-fold range generated by genetic modification (Doheny-Adams *et al*., 2012). Z007E0067 and Z007E0150 had similar stomatal density but Z007E0067 had slightly shorter stomata guard cell length compared to the Z007E0150 genotype (Table 1). However, Z007E0067 had consistently higher steady-state *g*_s_ and slower stomatal closure leading lower *WUE*_i_ which suggests the relatively small difference in guard cell length was not driving the differences in A_net_/*g*_s_. Ozeki et *al*., (2022) reported a correlation of guard cell length and stomata response time but the variation in stomatal length reported was ∼20 µm (4.75 fold larger) whereas the difference between Z007E0067 and Z007E0150 geontypes was just ∼4 μm (Table 1). Therefore, the differences in steady-state *g*_s_ and transient responses to environmental stimuli between the Z007E0067 and Z007E0150 genotypes likely stem from biochemical rather than anatomical or structural traits.

## Conclusion

Since natural variation in stomatal kinetics already exists it has potential to be integrated into crop species to improve *WUE*_i_ in dynamic environmental conditions. The results of this study suggest that this substantial variation in stomatal speed which was detectable by δ^13^C_leaf_ was directly inherited or recombined from one of the two parents of the NAM-Z007 family (B73 X CML333 RILs). These results demonstrate that one mechanism of enhanced *WUE*_i_ under dynamic conditions has promise for breeding for *WUE*_i_, especially for C_4_ crops because the CCM saturates at low *C*_i_ and is not diminished by low *g*_s_. C_4_ Δ^13^C theory has long predicted that variation in δ^13^C_leaf_ could be associated with *WUE*_i_ (Farquhar, 1983). These results confirm that variation in stomatal conductance can be screened by δ^13^C_leaf_ and linked to *WUE*_i_. However, we caution that screening with δ^13^C_leaf_ may not always predict *WUE*_i_ that is driven by stomatal kinetic traits. Variable QTL’s for δ^13^C_leaf_ from different populations just in *Z. mays* also suggests there are likely several factors impacting it which may associate to *WUE*_i_ or may not (Farquhar, 1983; Tcherkez *et al*., 2011; Gresset *et al*., 2014; Avramova *et al*., 2019). The reason this experiment was able to associate δ^13^C_leaf_ to fast-closing stomata and *WUE*_i_ is that it was in the context of genotypes with a known pedigree. More diverse surveys of δ^13^C_leaf_ and genetic variation could discover mechanisms that increase photosynthetic capacity as well as anatomical traits like stomatal density or restricted xylem vasculature. It is possible that the difficulty in interpreting δ^13^C_leaf_’s ability to predict *WUE*_i_ within diverse C_4_ populations in previous literature was because several traits could have influenced δ^13^C_leaf_ in opposing directions (Monneveux *et al*., 2007; Cabrera-Bosquet *et al*., 2009). Alternatively, post-photosynthetic fractions altering δ^13^C_leaf_ could obscure a link to *WUE*_i_ (Tcherkez *et al*., 2011). Looking to the future, screening for *WUE*_i_ with δ^13^C_leaf_ in structured populations combined with statistical genetic tools holds great potential for discovering and engineering mechanisms to improve *WUE*_i_.

## Supplemental Figures

**SFigure 1.**
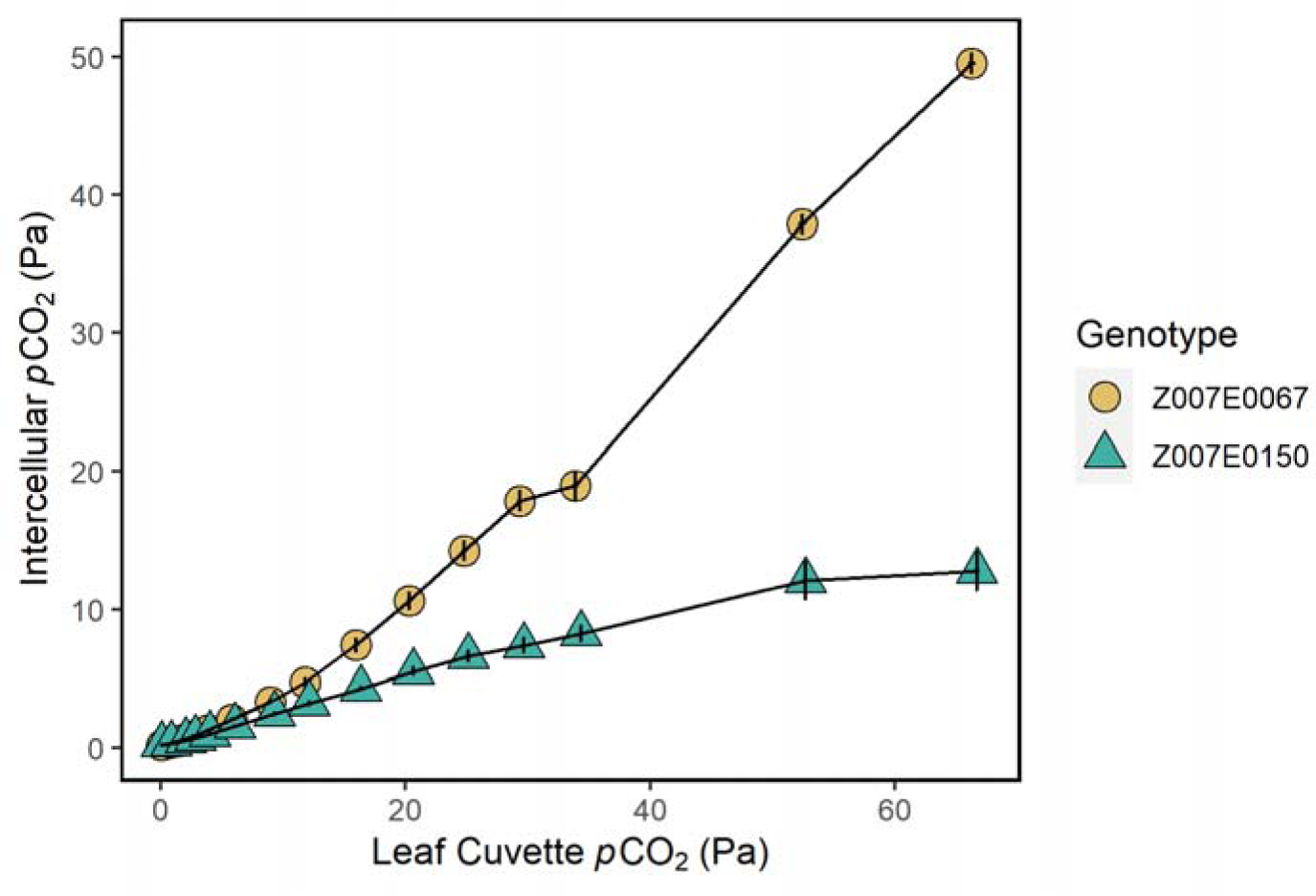
The internal leaf partial pressure of CO_2_ (C_i_) measured in response to *p*CO_2_ (Pa) surround the leaf in the *Z. mays* genotypes Z007E0067 and Z007E0150 with contrasting for carbon isotope composition (δ^13^C_leaf_). The measurements were at a leaf temperature of 30°C, 70% relative humidity, and Photosynthetic Photon Flux Density (PPFD) of 1,800 µmol m^-2^ s^-1^. Symbols in all panels are mean value ± SE (n=6). These data were collected from the measurements presented in Figure 1.

**SFigure 2.**
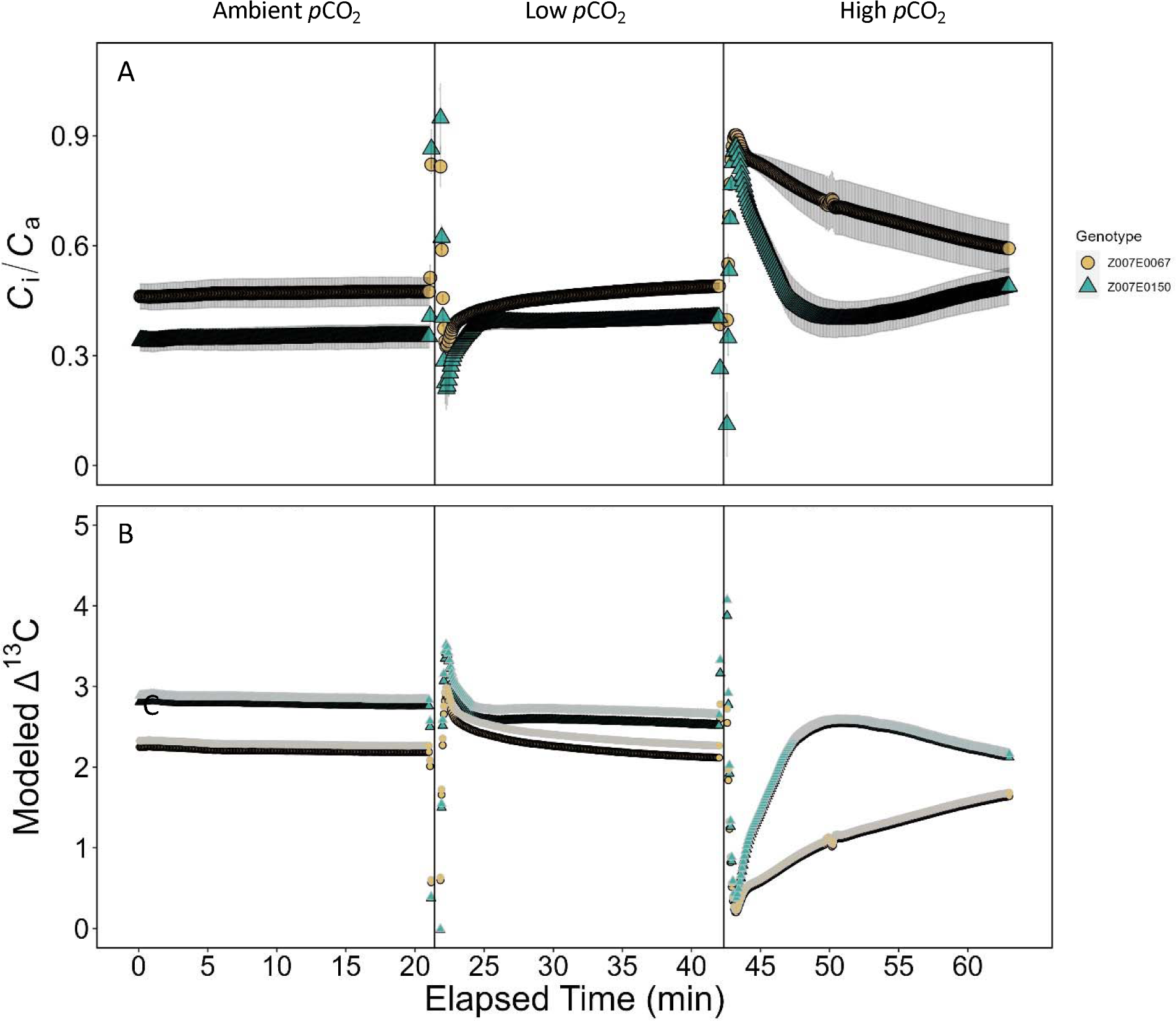
A time course response of *C*_i_/*C*_a_ and the modeled Δ^13^C resulting from differences in *C*_i_/*C*_a_ to step-changes in *p*CO2 in two *Z. mays* genotypes contrasting for carbon isotope composition (δ^13^C_leaf_). Ambient *p*CO_2_ (37 Pa); Low *p*CO_2_ (13.9 Pa), and High *p*CO_2_ (69.8 Pa). Each period of *p*CO_2_ lasted about 20 min. Data was collected on n=6 (Z007E0067) and n=5 (Z007E0150) and data points were collected every 5 sec. Symbols in all panels are mean value ± SE. In the points with black outlines a simplified modeled Δ^13^C is shown. Points with white outlines represent a modeled Δ^13^C with *g*_m_ conservatively modeled as 2 µmol m^-2^ s^-1^ Pa^-1^. The overlap suggests *g*_m_ is not the factor driving differences in Δ^13^C.

